# Tarantula phylogenomics: A robust phylogeny of multiple tarantula lineages inferred from transcriptome data sheds light on the prickly issue of urticating setae evolution

**DOI:** 10.1101/501262

**Authors:** Stephen Foley, Tim Lüddecke, Dong-Qiang Cheng, Henrik Krehenwinkel, Sven Künzel, Stuart J. Longhorn, Ingo Wendt, Volker von Wirth, Rene Tänzler, Miguel Vences, William H. Piel

## Abstract

Mygalomorph spiders of the family Theraphosidae, known to the broader public as tarantulas, are among the most recognizable arachnids on earth due to their large size and widespread distribution. Their use of urticating setae is a notable adaptation that has evolved exclusively in certain New World theraphosids. Thus far, the evolutionary history of Theraphosidae remains poorly understood; theraphosid systematics still largely relies on morphological datasets, which suffer from high degrees of homoplasy, and traditional targeted sequencing of preselected genes failed to provide strong support for supra-generic clades (i.e. particularly those broader than subfamilies). In this study, we provide the first robust phylogenetic hypothesis of theraphosid evolution inferred from transcriptome data. A core ortholog approach was used to generate a phylogeny from 2460 orthologous genes across 25 theraphosid genera, representing all of the major theraphosid subfamilies, except Selenogyrinae. For the first time our phylogeny recovers a monophyletic group that comprises the vast majority of New World theraphosid subfamilies including Aviculariinae and Theraphosinae. Concurrently, we provide additional evidence for the integrity of questionable subfamilies, such as Poecilotheriinae and Psalmopoeinae, and support the non-monophyly of Ischnocolinae. The deeper relationships between almost all subfamilies are confidently inferred for the first time. We also used our phylogeny in tandem with published morphological data to perform ancestral state analyses on urticating setae. This revealed that the evolution of this important defensive trait might be explained by three equally parsimonious scenarios.

## 1. Introduction

Theraphosidae is the largest family of mygalomorph spiders whose members are more commonly known as tarantulas or bird-eating spiders. Researchers have long been intrigued by these spiders due to both their large size and the plethora of adaptive traits they display. Tarantulas were painted by early naturalists as bird-eating monstrosities, and later as enemies of humans in horror movies, which together earned them a notorious reputation among the general public. However, interest in these spiders has boomed in recent decades (e.g. as in Teyssie 2015) and is still enthusiastically sustained among both hobbyists and researchers to this day.

At the time of writing, Theraphosidae contains 992 accepted species across 146 genera that are placed into 12 subfamilies by most authors (Kambas, 2018; World Spider Catalog, 2018). This species richness, combined with their widespread distribution and hence their diversity of habitats and ecological niches, is linked to a high variability of morphological and ecological adaptations. These adaptations have been the focus of a variety of studies in recent years. For example, the mechanisms and structures that are responsible for the vast array of theraphosid colorations (Hsiung et al., 2015), venom compositions (e.g. Rodríguez-Rios et al., 2017; Santana et al., 2017), adhesive capabilities (Pérez-Miles et al., 2017) as well as urticating setae (Bertani and Guadanucci, 2013) have all received a fair amount of attention. The latter represents a special feature in Theraphosidae. These setae are exclusive to certain Neotropical tarantulas, but generally rare in the animal kingdom, although some lepidopteran caterpillars have vaguely comparable analogues (e.g. Battisti et al., 2011). Urticating setae of theraphosids are barbed, microscopic setae that are usually localised on the opisthosoma (Teyssie, 2015). In a long-range defense mechanism loosely known as “hair flicking” or “bombardment” (Fig 1a), many Neotropical species will use their legs to slough off these setae from their opisthosoma and actively disperse them into the air when a threat is perceived (Cooke 1972; Bertani and Guadanucci, 2013). Other species opt for an active “direct contact” approach against potential threats, in which they seek to press them onto the target (after Cooke 1972; Bertani and Guadanucci, 2013; Perafán et al., 2016). Consequently, urticating setae have been known to induce painful symptoms in humans (Zilkens et al., 2012; McAnena et al., 2013). Contact with urticating setae may cause mammalian skin or eye irritations due to their harpoon-like structure, and inhalation has been known to result in respiratory problems (Schmidt, 2003; Bertani and Guadanucci, 2013; Teyssie, 2015). When directed against predators, these defenses may provide the tarantula an opportunity to escape from predation (Cooke 1972). Additionally, some species also utilize urticating setae in a passively defensive manner, where they are attached to silk that covers egg-sacs or forms molting webs. This has been shown to be effective against invertebrates, in particular countering the larvae of parasitic flies (i.e. phorids) or ants (Marshall and Uetz, 1990a; Bertani and Guadanucci, 2013).

**Fig. 1:**
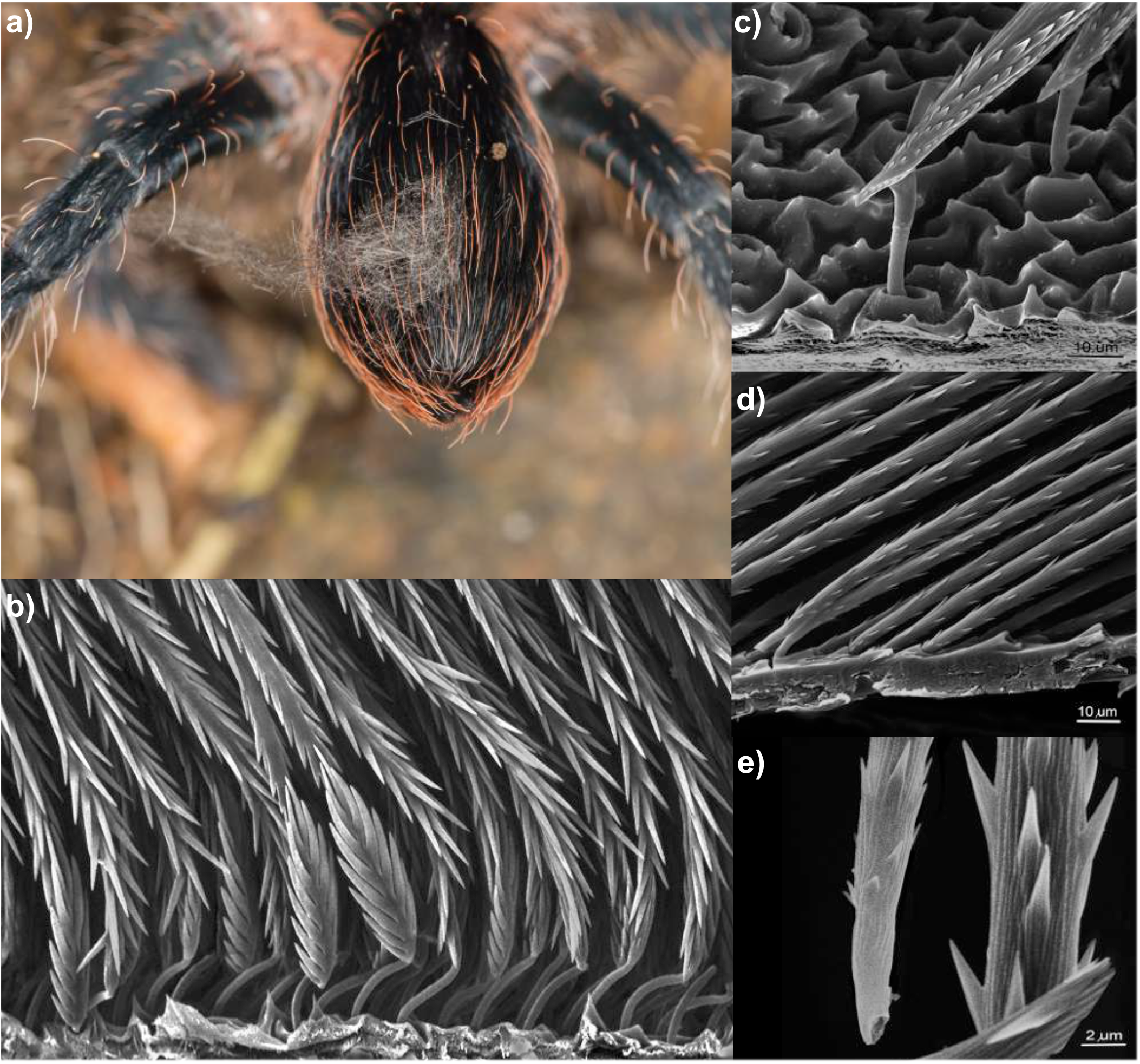
Urticating setae as a means of defense in tarantulas. In most cases urticating setae are sloughed off by the spider from its dorsal or lateral opisthosoma and projected into the air, as given in a) (photography: Martin Hüsser). The barbed and harpoon-like microstructure of urticating setae in selected tarantulas is highlighted in b) □ e) (photographs courtesy of Rainer Foelix): b) A dense assemblage of type I & III setae in *Brachypelma* sp. from the opisthosoma, c) type II setae from *Avicularia* sp. from the opisthosoma. d) type V from *Ephebopus* from the palpal femur and e) more closely detailed.

Seven different types of urticating setae have been described and they are distinguished by characters such as their microstructure, localization, or release mechanism (Bertani and Guadanucci, 2013; Perafán et al., 2016). The types I, III, IV, VI and VII occur in members of the Theraphosinae subfamily (Fig. 1b), whereas type II is exclusive to Aviculariinae (Fig. 1c). Certain types are even genus-specific, such as type V for *Ephebopus* (Fig. 1d-e), type VI for *Hemirrhagus* and type VII for *Kankuamo* (Bertani and Guadanucci, 2013; Foelix, Bastian and Erb 2009; Marshall and Uetz, 1990b; Mendoza & Francke 2018; Perafán et al., 2016). This conservation and specificity to certain groups of tarantulas has led to their widespread use as diagnostic taxonomic characters (e.g. Schmidt, 2003; Turner et al., 2018).

Given the species richness, charismatic nature and the breadth of scientific work on their adaptations, it is rather surprising and unfortunate that the taxonomic status and evolutionary history of theraphosids in general remains poorly understood. Most previous works on theraphosid evolution and systematics have been based exclusively upon the analysis of morphological characters (e.g. West et al., 2008; West and Nunn, 2010; Guadanucci, 2014; Fukushima and Bertani, 2017), which have often yielded inconsistent results. Subsequent studies recognized the limitations of purely morphological analyses due to high degrees of homoplasy (e.g. Ortiz et al., 2018). The use of molecular characters to study theraphosid evolution is relatively recent (e.g. Longhorn et al., 2007; Petersen et al., 2007; Hamilton et al., 2016; Ortiz and Francke, 2016; Turner et al., 2018; Mendoza and Francke, 2017; Ortiz et al., 2018; Hüsser, 2018), and a first comprehensive analysis of major theraphosid lineages was able to support the monophyly of Theraphosidae itself as well as of many subfamilies within (Lüddecke et al., 2018). Further, it provided molecular evidence for the validity of certain subfamilies that were previously uncertain, and therefore highlighted the potential for molecular data to resolve the theraphosid phylogeny. Despite this first study targeting a handful of commonly used nuclear and mitochondrial genes, the deeper phylogenetic relationships of theraphosid lineages were poorly supported and remained uncertain.

Well-supported phylogenies are key to making meaningful inferences about evolution, phylogeography, adaptation, and beyond. In this study, we aim to address this deficiency by providing a phylogenomic perspective on theraphosid evolution with a robust backbone inferred from an extensive molecular dataset. We sequenced 29 transcriptomes from a comprehensive range of disparate species, and combined these with 4 other published transcriptomes to generate a phylogenetic hypothesis that includes representatives of all major subgroups within the family. We analysed several subsets of orthologous genes (OGs) under varying measures of stringency with the intent to (i) resolve deeper clades within the phylogeny of Theraphosidae, (ii) test whether strongly supported nodes will be recovered in congruence with previous morphological and molecular studies as a means to assess their validity, and to (iii) reconstruct the ancestral states of urticating setae, an excellent model for clade-specific traits with important biological functions. Further, this phylogeny will serve as the groundwork for future evolutionary studies to examine other morphological adaptations in Theraphosidae, as well as other aspects such as evolution of venom components and biogeographical analyses.

## 2. Materials and Methods

### 2.1 Sampling, RNA Extraction and Sequencing

Detailed information regarding the particulars of each sample is available in supplementary Table S1, along with voucher deposition numbers. Sample material for this study was obtained either from the pet trade, private breeders, or collected in the field. Each specimen was identified using published identification keys, at least to genus level (Schmidt, 2003; Teyssie, 2015). Whenever possible, whole bodies of each specimen were sampled for this study, but in some cases only autotomized legs were available. Samples were either stored in RNAlater (coded as SFx in Table S1), or were pre-frozen in liquid nitrogen prior to freezing at −80°C (coded as TPx or RFx in Table S1). RNA extractions were performed either via the TRIzol total RNA extraction protocol or traditional phenol chloroform extractions (Simms et al., 1993). Purified RNA was sent to the Max Planck Institute in Plön, Germany, or to commercial sequencing companies who prepared cDNA libraries, and sequenced the samples using Illumina HiSeq and NextSeq technologies. These transcriptomes were complemented with publicly available data for four additional species acquired from the NCBI SRA database (http://www.ncbi.nlm.nih.gov/sra). We endeavoured to confirm that any variance from using different sequencing technologies and transcripts from different tissues was minimized by including pairs of conspecific (2x *Neoholothele incei* and 2x *Poecilotheria vittata*) and congeneric (2x *Caribena*, 2x *Psalmopoeus*, 2x *Pterinochilus* and 2x *Cyriopagopus*) taxa under the expectation that they would group together. Both the sequencing method and tissue sample for each member per pair differed from the other. All newly obtained transcriptome raw data were submitted to SRA under Bioproject number (to be added upon manuscript acceptance). Assembly data for newly sequenced transcriptomes is available in supplementary Table S2.

### 2.2 Core Ortholog Analysis Pipeline for Generating the Phylogeny

Transcriptomes were assembled using Trinity (Grabherr et al., 2011). Prediction of protein coding regions was performed in TransDecoder (Haas et al., 2013). A core ortholog approach based on Garrison et al. (2016) and modified by Cheng and Piel (2018) was used for putative ortholog selection, and aims to select only the genes that are orthologous across our taxon set. Following from Cheng and Piel (2018), a core ortholog set was derived from transcripts of *Damon variegatus, Acanthoscurria geniculata, Dolomedes triton, Ero leonina, Hypochilus pococki, Leucauge venusta, Liphistius malayanus, Megahexura fulva, Neoscona arabesca, Stegodyphus mimosarum*, and *Uloborus* sp., all of which are publicly available from SRA, and a set of 4,446 spider-specific profile hidden Markov models (pHMMs) was then built following from Cheng and Piel (2018). HaMStR v.13.1 (Ebersberger et al., 2009) was used to infer orthology between the pHMMs and sequences from each taxon analysed here (see Supp. Tab. S1). Groups of orthologous genes (OGs) corresponding to the pHMMs were subsequently pooled and subjected to several rounds of filtering: Sequences shorter than 75 amino acids and OGs sampled for fewer than 17 specimens were discarded. MAFFT (Katoh et al., 2005) was used to align each OG, with the settings “auto”, “localpair” and “maxiterate 1000” selected. Alignments were trimmed by ALISCORE (Misof and Misof, 2009; Kück et al., 2010). Ambiguous regions were excised in ALICUT (Kück, 2009), and Infoalign (Rice et al., 2000) was used to build consensus sequences for each alignment. In an effort to remove paralogs, sequences that were far from the consensus (i.e. with an infoalign *change* value between the sequence and consensus exceeding 75) were removed. Sequences with >9 gaps flanking a region with 20 of fewer mistranslated amino acids were excluded, as were alignment columns with <4 non-gap characters. Any sequences that were shorter than 75 amino acids after these filtering steps were also removed. Finally, sequences that did not overlap with all other sequences in the alignment by at least 20 amino acids were removed, and OGs sampled for fewer than 17 taxa were discarded. This yielded our “full set” of OGs, which constitute our first dataset (matrix 1).

Following Cheng and Piel (2018), data for four additional OG matrices with varying degrees of conservation was derived via matrix reduction and optimization. These resulted from custom scripts (Cheng and Piel, 2018). The matrix 1 dataset was sorted first by gene occupancy, and then by gene length. OGs composing matrix 2 (“first reduce”) were obtained by only retaining the larger half of the sorted matrix 1 dataset. OGs composing matrix 3 (“second reduce”) were obtained by only retaining the larger half of matrix 2 dataset. OGs composing matrix 4 were selected using BaCoCa (Kück and Struck, 2014), which optimised the full matrix 1 dataset to retain only 50% composed of the most phylogenetically informative sites. OGs for matrix 5 were selected using MARE (Meyer et al., 2011), which assessed the matrix 1 dataset partitions, provided a measure of tree-likeness for each gene, and finally optimized the matrix for information content with an alpha value of 5. FASconCAT (Kück and Meusemann, 2010) was used to concatenate each of the five OG datasets to yield the five data matrices. Metrics for all matrices, including numbers of OGs and amino acids per dataset, are available in Table 1.

**Table 1:** Metrics of the five OG matrices, including the number of OGs per matrix, corresponding amino acid numbers (AAs), the percentage of missing data, and the log likelihood of the best tree for each matrix.

| ANALYSIS | #OGs | #AAs | % MISSING | LOG LIKELIHOOD |
| --- | --- | --- | --- | --- |
| Matrix 1: All genes | 2460 | 1096124 | 16.85% | -9703757.281168 |
| Matrix 2: First reduce | 1230 | 628946 | 7.92% | -5534973.862171 |
| Matrix 3: Second reduce | 615 | 400656 | 5.36% | -2883038.747349 |
| Matrix 4: BaCoCa | 1230 | 712868 | 14.34% | -6099109.419374 |
| Matrix 5: MARE | 1548 | 723749 | 9.80% | -5554777.432207 |
| ASTRAL | 2460 | - | - | - |

ExaML v.3.0.2 (Kozlov et al., 2015) was used for maximum likelihood searches, and optimal trees were calculated for each matrix, and optimal models for amino acid substitutions were determined using the AUTO command. As in Cheng and Piel (2018), the gamma parameter was estimated using a model of rate heterogeneity with four discrete rates. Parsimony trees constructed by RAxML v.8.2.11 were used to initiate the searches. Bootstrapping was performed in ExaML v.3.0.2 with the same parameters as above, with 200 replicates generated per dataset.

ASTRAL multispecies coalescent analysis (Mirabab et al., 2014) was also used as a sixth approach, but as it is particularly sensitive to paralogy, we implemented an additional method of ortholog filtering. Rooted gene trees for each OG were generated by RAxML v.8.2.11 (Stamatakis, 2014), and these trees were pruned to further excise putatively paralogous sequences using a custom script. This script, a modification from the original pipeline, uses the rooted trees to group each taxon into bins based on the distribution of root-to-tip lengths. The script prunes small subsets of taxa with root-to-tip lengths that greatly exceed the bulk of other taxa on the expectation that truly orthologous genes approximate a molecular clock. These pruned gene trees served as inputs for ASTRAL, which estimated the coalescent species tree.

### 2.3 Gene Jackknifing

Analysis of large phylogenomic data sets can result in maximum support for nodes even if the actual support is rather spurious in the data. In such cases, wrong topologies might be inferred with high confidence due to the effects of even very minor cross-contamination, paralogy, or alignment errors remaining in the data set despite stringent filtering (Philippe et al., 2011). To better understand if any of the nodes in our phylogenomic analyses may be affected by such phenomena, gene jackknifing was carried out following Irissari et al. (2017). We generated 100 gene jackknife replicates for concatenated OG subsample alignments of 10,0, 100,000, 500,000 and 1,000,000 amino acids in length. Maximum likelihood trees for each replicate were estimated in RAxML v.8.2.11 and majority rule consensus trees for each of the four differently sized jackknife matrices were generated by concatenating their respective 100 replicate trees into a single file each. From these files, we examined selected nodes in our phylogeny to see whether they stabilized with high support values with smaller data subsets (strong phylogenetic signal), or whether they required a larger proportion of our original data (limited phylogenetic signal).

### 2.4 Ancestral State Reconstruction of Urticating Setae

A multistate character matrix detailing the presence and absence of different urticating setae types among New World species in our phylogeny was constructed (available in Table S3) in accordance with the literature (e.g. Bertani and Guadanucci, 2013; Hüsser, 2018). We assigned urticating setae of types I, III and IV to different taxa from the subfamily Theraphosinae (after Bertani and Guadanucci, 2013), type II to Aviculariinae (Fukushima and Bertani 2017), and type V solely to the genus *Ephebopus* (Foelix et al., 2009; Marshall and Uetz, 1990b). Urticating setae types VI and VII, each known from only *Hemirrhagus* and *Kankuamo* respectively, were not included in this analysis due to a lack of sample material. Ancestral state reconstruction was performed in Mesquite v3.51 (Maddison and Maddison, 2018) using default parameters with this matrix. Character states were treated as unordered.

## 3. Results

### 3.1 Metrics of Tarantula Phylogenomics

New transcriptomes were generated from 28 samples of 26 theraphosid species, plus another from the family Dipluridae. Combined with existing transcriptomic data from four additional species, we generated five OG matrices that ranged in size from 615 to 2460 OGs, with a total length of 1,096,124 amino acids for the largest “all genes” matrix of all OGs combined (Table 1). Additional derivative matrices 1^st^ reduce and 2^nd^ reduce, which decreased the matrix size by half each time through removal of genes with the greatest proportion of missing data, improved completion from 16.85% missing data in the initial matrix down only to 7.92% in 1230 OGs and 5.36% in 615 OGs respectively. Matrix 4 (BaCoCa) had 14.43% missing data despite containing the same number of OGs as matrix 2 (1203). Matrix 5 (MARE) had 9.80% missing data and contained a large subset of OGs (1548). All conspecifics and congenerics found each other. The number of OGs for each taxon that matched to the initial spider-specific set of 4,446 pHMMs (“SPIDs”) ranged from 2611 (*Psalmopoeus irminia*) to 3578 (*Thrigmopoeus* sp.). The total numbers of OGs per taxon in the “full matrix” (matrix 1) ranged from 1691 (*Linothele* sp.) to 2348 (*Thrigmopoeus* sp.). A full list of SPIDs, total OGs in matrix 1, and assembly data for all newly sequenced transcriptomes are available in Table S2.

Bootstrap values of 100% were recovered for all but two nodes (given in Fig. 2) across all six of our phylogenomic analyses with maximum likelihood. Overall, our phylogeny includes 29 theraphosid species, representing 25 genera and 11 subfamilies. Members of two other mygalomorph families, Dipluridae (*Linothele* sp.) and Nemesiidae (*Damarchus* sp.) were included as outgroups to root the phylogenetic trees.

**Fig. 2:**
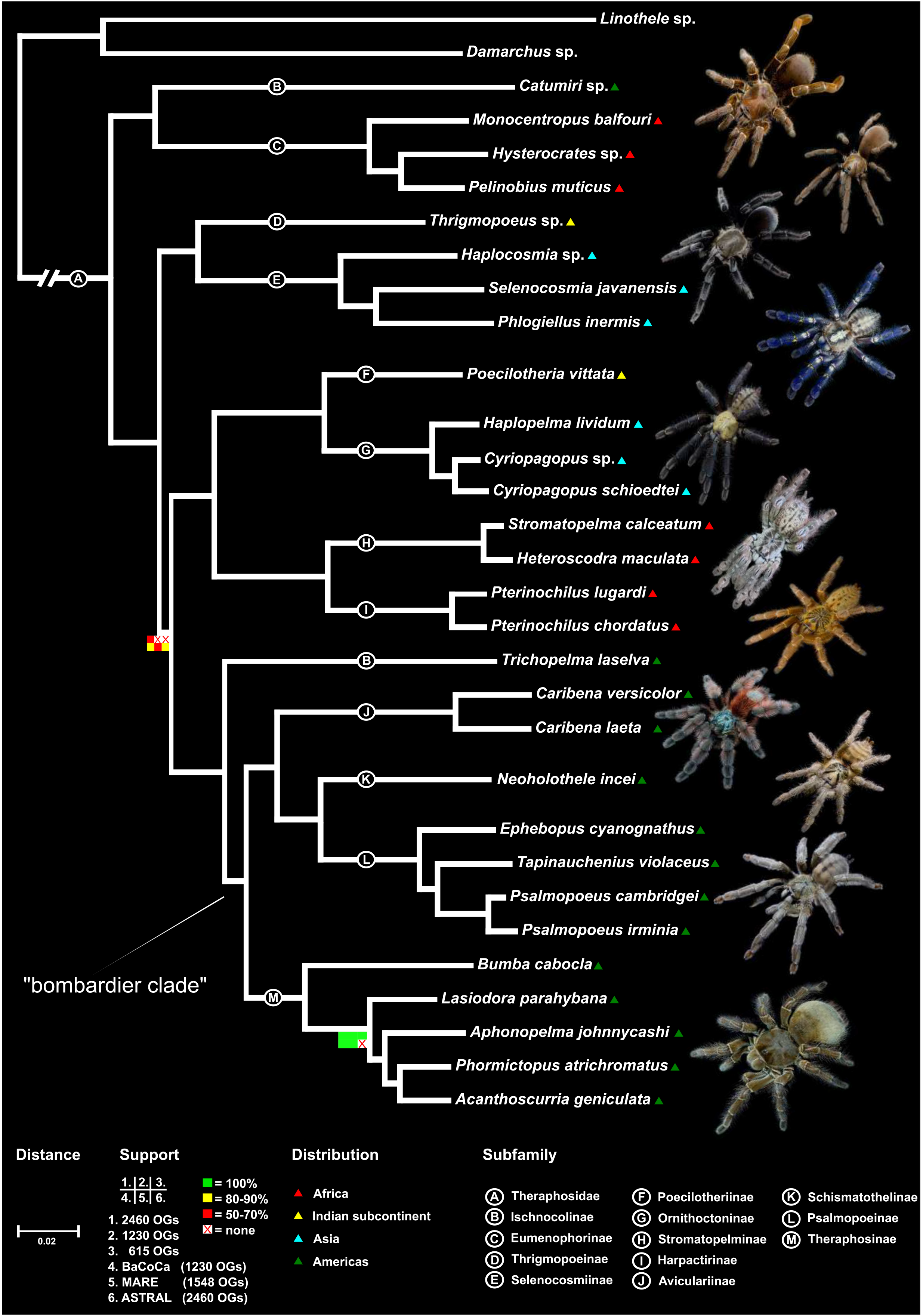
Summary result of our maximum likelihood phylogenomic tree of Theraphosidae generated using ExaML. The tree obtained from matrix 1 is shown to serve as the background tree, upon which node support values for each of the other five analyses are superimposed. Subfamilial designations, support values, distances and geographic range are provided. Bootstrap support values are 100% unless otherwise indicated. Images depict representative taxa of included subfamilies. Photographs courtesy of Bastian Rast.

### 3.2 The Deep “Tarantula Tree of Life “ inferred from Transcriptome Data

Our combined results (Fig. 2) recovered Theraphosidae as monophyletic. The first clade in our phylogeny to branch, splitting from remaining theraphosids, consists of African representatives of the Eumenophorinae subfamily (*Hysterocrates* sp., *Monocentropus balfouri* and *Pelinobius muticus*), and an American member of the Ischnocolinae (*Catumiri* sp.). Branching off next as sister to the remaining theraphosids is a clade composed of Indian Thrigmopoeinae (represented by a single taxon, *Thrigmopoeus* sp.), and Selenocosmiinae from across many parts of Asia and Australasia (*Haplocosmia* sp., *Phlogiellus inermis* and *Selenocosmia javanensis*).

The next node is the only one that did not receive consistently high bootstrap support, which includes a first clade branching off that consists of four subfamilies and two subclades. This first subclade includes two African subfamilies — the Stromatopelminae (*Heteroscodra maculata* and *Stromatopelma calceatum*) that are placed sister to Harpactirinae (*Pterinochilus chordatus* and *Pterinochilus lugardi)*. The second subclade consists of Poecilotheriinae (actually from two replicates of *Poecilotheria vittata* generated from different individuals), which are endemic to the Indian subcontinent, and their sister group Ornithoctoninae (*Haplopelma lividum, Cyriopagopus* sp. and *Cyriopagopus schioedtei*), members of which are found throughout Southeast Asia and parts of China. The second clade exclusively harbors representatives of five subfamilies of Theraphosidae from the Americas: Splitting first is a member of the Ischnocolinae (*Trichopelma laselva*). The rest of this American clade is further divided in two main subclades. The first of these includes only members of the subfamily Theraphosinae (*Acanthoscurria geniculata, Aphonopelma johnnycashi, Bumba cabocla, Lasiodora parahybana* and *Phormictopus atrichomatus*). The second clade consists of three subfamilies, with Aviculariinae (*Caribena versicolor* and *Caribena laeta*) as sister group of both Schismatothelinae (two replicates of *Neoholothele inceî*) and Psalmopoeinae, with *Ephebopus* (*Ephebopus cyanognathus*), as sister to the remaining species (*Psalmopoeus cambridgei, Psalmopoeus irminia* and *Tapinauchenius violaceus*).

With the exception of the “ischnocoline” species *Catumiri* sp., we have recovered all New World theraphosids in our phylogeny as a monophyletic group. The subfamilies Eumenophorinae, Harpactirinae, Stromatopelminae, Theraphosinae, Selenocosmiinae, Ornithoctoninae, Psalmopoeinae and Aviculariinae, included in our analyses with at least two species each, were recovered as monophyletic with 100% support across all analyses. The monophyly of Thrigmopoeinae, Poecilotheriinae and Schismatothelinae cannot be confirmed due to the availability of only a single species for each subfamily, and the known-to-be paraphyletic Ischnocolinae were recovered as such.

Contrary to prior studies, our phylogeny is distinguished by high bootstrap support values across almost all nodes, with all but two nodes recovering 100% support across all six analyses. The placement of *Lasiodora parahybana* as sister taxon to three other Theraphosinae (*Aphonopelma johnnycashi, Acanthoscurria geniculata* and *Phormictopus atrichomatus*) received 100% bootstrap support in most analyses. *Aphonopelma johnnycashi* was absent from matrix 5 (MARE), although adjacent nodes nonetheless received 100% bootstrap support. In the ASTRAL topology, *Aphonopelma johnnycashi* was instead recovered as the sister taxon to *Lasiodora parahybana, Acanthoscuria geniculata* and *Phormictopus atrichomatus* and recieved 100% bootstrap support. The node defining the clade placed sister to Selenocosmiinae + Thrigmopoeinae received bootstrap values of 55% (matrix 1), 90% (BaCoCa), 69% (MARE) and 82% (ASTRAL). Matrices 2 and 3 recovered an alternative topology where Selenocosmiinae + Thrigmopoeinae branched alongside Poecilotheriinae / Ornithoctoninae / Harpactirinae / Stromatopelminae, with bootstrap values of 81% and 56% respectively.

Gene jackknifing confirmed high support (Gene Jackknife Proportions, GJP > 70%) already with gene sampling at 10 Kbp, and GJP = 99-100% with 100 Kbp (Supplementary Table S4). This suggests an extraordinarily strong phylogenetic signal in our data set. As the only two exceptions, (i) the node joining *Lasiodora parahybana + Aphonopelma johnnycashi* received GJP < 50% at 10 and 100 Kbp and only stabilized (GJP = 71% and 100%) with 500 Kbp and 1 Mbp), and (ii) the node defining the clade sister to *Thrigmopoeus* + Selenocosminae (which also received poor bootstrap support in several analyses; Fig. 2) received poor GJP around or below 50% up to inclusion of 500 Kbp, and only started stabilizing at 72% with 1 Mbp data sets.

### 3.3 Three possible Scenarios for the Evolution of Urticating Setae

We identified a clade in our phylogeny that includes the members of four New World theraphosid subfamilies: Aviculariinae, Psalmopoeinae (including *Ephebopus*), Schismatothelinae and Theraphosinae. All of the species in our phylogeny that possess urticating setae, and who have been observed to engage in hair-bombardment behavior, are included in these subfamilies. Hence, we will refer to this clade as the “bombardier clade”. While urticating setae types I, III and IV are present in Theraphosinae, one of the two clades arising from the earliest internal split, type II setae are only found in some members of its other clade, corresponding to the Aviculariinae. Schismatothelinae lack urticating setae, as do the Psalmopoeinae genera (e.g. *Psalmopoeus* and *Tapinauchenius*). However, the psalmopoeine *Ephebopus* (represented in our phylogeny by *Ephebopus cyanognathus*) is the only genus with type V urticating setae. This array of urticating setae types found in our sampled taxa are coded in supplementary Table S3. Our ancestral state reconstruction of urticating setae evolution within the bombardier clade fits the characters onto our topology with 7 steps. The result (Fig. 3a) illustrates the different urticating setae types traced through the bombardier clade. Consistent with previous studies, our Fig. 3b presents a “multiple-gain” scenario where urticating setae have independently evolved at least three times during the evolution of Theraphosidae. However, Fig. 3c presents an equally parsimonious “multiple-loss” alternative, whereby urticating setae evolved a single time and were then subsequently lost twice, once in Schismatothelinae and again in non-*Ephebopus* Psalmopoeine. Lastly, Fig. 3d presents another equally parsimonious “gain-loss-gain” scenario of urticating setae evolution, with setae being lost when Aviculariinae and Schismatothelinae diverged, but subsequently regained in *Ephebopus*.

**Fig. 3:**
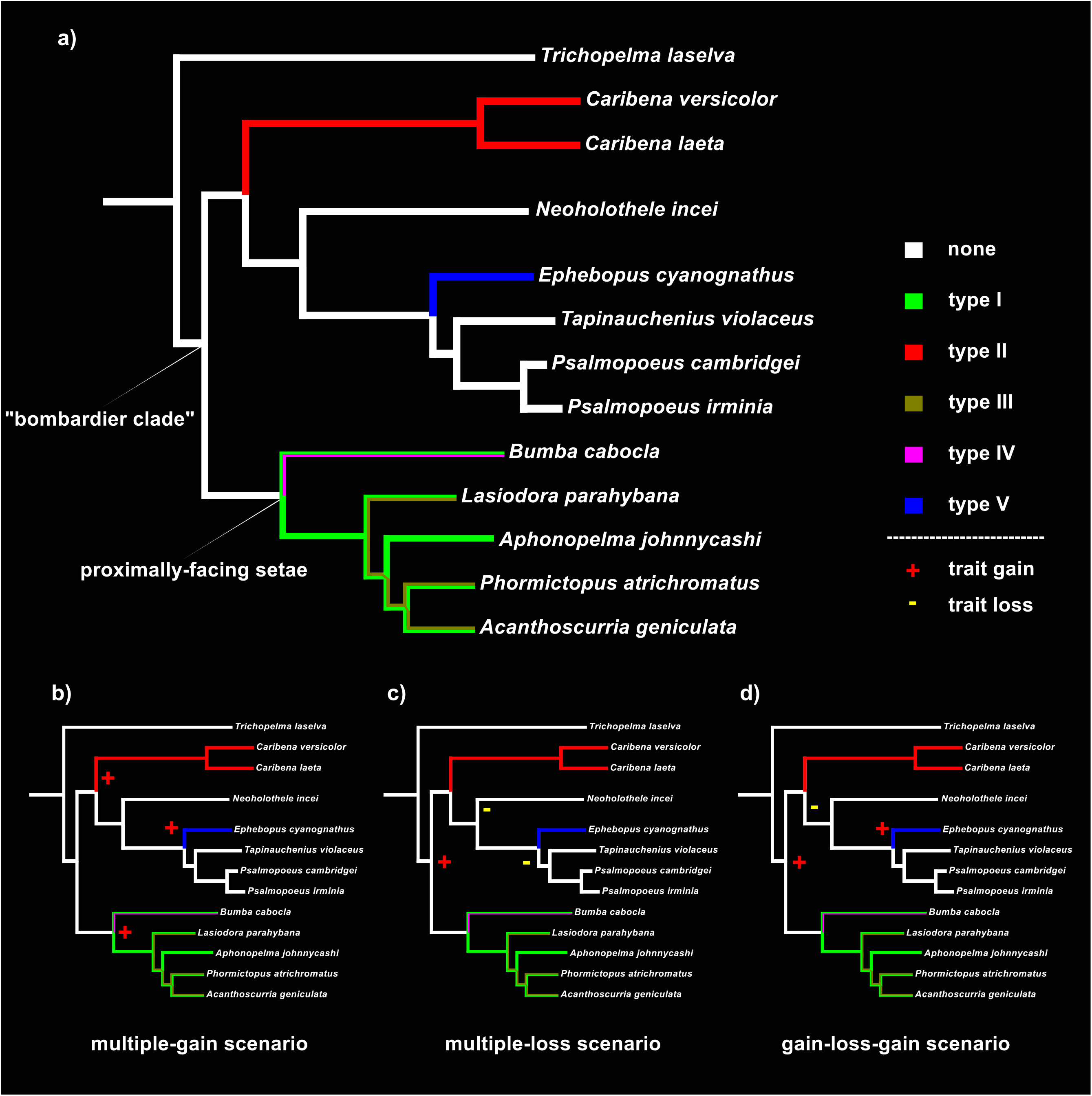
Overview of urticating setae evolution among theraphosids, with three equally parsimonious hypotheses. a) The ancestral state reconstruction of different urticating setae types among our “bombardier” clade, and the shared ancestry of “proximally-facing” bristles types I, III and IV. b) Prevailing hypothesis whereby urticating setae evolved three separate times independently in Theraphosinae, Aviculariinae and *Ephebopus* (Psalmopoeinae). c) Hypothesis of a single origin for urticating setae with one origin leading to types I, II, III and IV in Theraphosinae and Aviculariinae, then subsequently being lost in both Schismatothelinae and then Psalmopoeinae (after the divergence of *Ephebopus*). d) Hypothesis of two potential independent origins for urticating setae, with one origin again, then subsequently being lost when the common ancestor for Schismatothelinae and Psalmopoinae diverged from Aviculariinae, before re-emerging in Psalmopoeinae via *Ephebopus* as type V setae.

## 4. Discussion

### 4.1 A First Phylogenomic Overview of Theraphosidae

This study marks the first time in which phylogenomic data has been applied to study the evolution of theraphosid spiders. We opted for a transcriptomic approach because low to moderate coverage can be an efficient means to obtain sequences of large numbers of protein-coding ortholog genes from the nuclear genome (e.g. Irisarri et al., 2017) and therefore constitutes a promising method for resolving deeper nodes in the tarantula tree of life. In our case, all but two nodes were recovered consistently with 100% bootstrap support in all phylogenomic analyses. As a result, we confidently infer for the first time many of the relationships between subfamilies and their placements within the overall theraphosid tree. Although both the number of SPIDs and total number of OGs recovered in matrix 1 was generally greater in taxa from which whole body samples were taken, all conspecifics and congenerics found each other.

Consistent with previous molecular studies, such as Lüddecke et al. (2018), we recovered Eumenophorinae as an early branching subfamily. However, we also recovered *Catumiri* sp. as its sister-group — a member of the taxonomically unresolved Ischnocolinae and not included in Lüddecke et al. (2018). It has been suggested that members of Ischnocolinae may represent some of the most primitive members of Theraphosidae (after Raven, 1985; Schmidt, 2003; Guadanucci 2014), which is at least partially supported by our results given that we found *Catumiri* placed near the base of our phylogenetic tree.

The next earliest split leads to another clade composed of two Asian subfamilies: Selenocosmiinae and Thrigmopoeinae. The former is found in Southeast Asia (plus some Indian subcontinent and China) and Australasia, while the latter is found exclusively in India, and has received little attention in previous comprehensive studies of theraphosids. Yet, Thrigmopoeinae has a long history of taxonomic revisions and many synonyms exist in this group (e.g. Prasanth and Sunil Jose, 2014; Sanap and Mirza, 2014; Sankaran and Sebastian, 2018). In previous morphological analyses (e.g. Raven, 1985; Guadanucci, 2014), Ornithoctoninae were found to be the closest relatives to Thrigmopoeinae. The molecular analysis of Lüddecke et al. (2018) was also in line with this relationship, placing Thrigmopoeinae and Poecilotheriinae together as a clade sister to Ornithoctoninae. This hypothesis was further supported from the perspective of biogeography, as the placement of Thrigmopoeinae with Poecilotheriinae results in a clade of taxa endemic to the Indian subcontinent. Contrary to all previously mentioned studies, our phylogenomic hypothesis supports a closer relationship of Thrigmopoeinae with Selenocosmiinae — which also has some representatives from Indian subcontinent as part of this subfamily’s broad geographic distribution, and rejects a placement of Thrigmopoeinae near Ornithoctoninae — which are more exclusively restricted to Southeast Asia, plus parts of China.

Next is a clade that contains four subfamilies that form two clades. The two African subfamilies (Harpactirinae and Stromatopelminae) form the first and two Asian subfamilies (Ornithoctoninae and Poecilotheriinae) the second clade. Although the molecular results of Lüddecke et al. (2018) already recovered the close relationship of Harpactirinae and Stromatopelminae, this connection has been controversial in the past. With morphological data, Stromatopelminae have historically been placed with Eumenophorinae (Raven, 1985), or inside Aviculariinae (West et al., 2008; Guadanucci, 2014; Fukushima and Bertani, 2017). Harpactirinae were otherwise thought to be more closely related to Theraphosinae and Aviculariinae (Raven, 1985), or even Schismatothelinae (Guadanucci, 2014). However, these hypotheses are rejected by our phylogenomic analysis, and we instead validate the close relationship of Harpactirinae and Stromatopelminae, and is supported by some morphological data (Gallon, 2003). Regarding the sister clade, the placement of Poecilotheriinae in the theraphosid tree has long been the subject of debate, with its members having most recently been included in Selenocosmiinae (e.g. Kambas 2018, following Raven, 1985). The closer relationship between Poecilotheriinae and Ornithoctoninae was only recently revealed by molecular data in Lüddecke et al. (2018), and is here further supported by our far more comprehensive transcriptomic approach.

A further major clade identified in our study consists exclusively of taxa from the Americas (our “bombardier” clade), and includes another member of Ischnocolinae, the Central American *Trichopelma laselva*, that branches off as sister-group to the rest. The clade is then divided into the subfamily Theraphosinae, and three other subfamilies Aviculariinae, Psalmopoeinae and Schismatothelinae that together form another clade. The placement of Theraphosinae as one of the more derived subfamilies is remarkable since it differs from a wide range of previously published studies (e.g. Raven, 1985; Guadanucci, 2014; Lüddecke et al., 2018), all of which placed Theraphosinae more basally, although that alternative placement did not receive strong support in the molecular study of Lüddecke et al. (2018). Our phylogenomic results instead indicate that Theraphosinae is one of the most recent subfamilies. Biogeographically this is of relevance, as our analysis suggests a monophyletic group of all of the New World taxa in our phylogeny, except for *Catumiri* sp. Given that members of the genus *Catumiri*, like most Ischnocolinae (if indeed this is where they should be placed), are characterised by the absence of many diagnostic characters such as urticating setae or stridulatory bristles (Schmidt, 2003), *Catumiri* sp. may represent an ancient biogeographic relic and the novel placement indicated by our new results certainly warrants further study. The placements of Aviculariinae, Psalmopoeinae and Schismatothelinae together as other relatively derived subfamilies is consistent with the multigene results of Lüddecke et al. (2018), which also recovered a corresponding clade for these, inline with other results from single genetic fragments (Turner et al., 2018; Hüsser, 2018).

### 4.2 Implications for Subfamilial Validity in Theraphosidae

While Theraphosidae itself and most of its well-established subfamilies, namely Aviculariinae, Eumenophorinae, Harpactirinae, Ornithoctoninae, Selenocosmiinae, Stromatopelminae and Theraphosinae, have emerged as monophyletic from our analysis, there are nevertheless some novel results that encourage us to reconsider the validity of certain subfamilies.

The subfamily of Poecilotheriinae from India and Sri Lanka has long been one of the most controversial from a taxonomic perspective. It was initially established as a unique lineage for just two genera including the genus *Poecilotheria*, which was later transferred to the Selenocosmiinae subfamily primarily due to the morphology of their stridulatory organs (e.g. Raven, 1985). However, this placement has always been controversial (e.g. Schmidt 1995, who argued instead for a monotypic Poecilotheriinae), and no universally accepted consensus surrounding the placement of *Poecilotheria* has so far been reached by morphological analysis. Molecular studies, on the other hand, recently demonstrated that the genus is genetically very distinct from Selenocosmiinae, and instead seems to share a more recent common ancestor with Ornithoctoninae. It has therefore been suggested that the placement of *Poecilotheria* inside Selenocosmiinae is not justified, and that the validity of Poecilotheriinae as a subfamily should be considered (Lüddecke et al., 2018). In agreement with this, our phylogenomic data also supports the distinction of *Poecilotheria* from Selenocosmiinae and instead supports their closer relationship with Ornithoctoninae as previously suggested by much smaller molecular datasets (Lüddecke et al., 2018; Turner et al., 2018). However, *Poecilotheria* differs from Ornithoctoninae not only genetically but further in respect to morphology: They lack plumose bristles on the retrolateral surface of chelicerae, which are diagnostic for Ornithoctoninae (Von Wirth and Striffler, 2005; West and Nunn, 2010). Given this genetic and phenotypic distinction of *Poecilotheria*, we follow Schmidt (1995) to emphasize in agreement with Lüddecke et al. (2018) that the unique subfamilial status Poecilotheriinae Simon 1892 stat. rev. should be adopted, with *Poecilotheria* Simon, 1885 as its type genus. This group will require a more detailed revision, ideally based on morphological as well as molecular data from a larger set of species, in the near future.

Among all theraphosid subfamilies, the Ischnocolinae are recognized as the taxonomically most problematic group. These are mostly relatively small spiders, often referred to as “dwarf tarantulas”, have the widest distribution among all theraphosids, with species being found on many continents, such as the Americas, Asia, Africa and even Europe. Originally, Ischnocolinae was described by Simon 1892 with all tarsi divided being the main diagnostic morphological character. However, it later became apparent that this trait is not unique to ischnocolines (e.g. Schmidt, 2003). In addition, other key tarantula traits such as urticating setae, stridulatory organs or basally connected spermathecae were never found in Ischnocolinae, resulting in a paucity of other informative characters (after Raven 1985; Schmidt, 2003; Guadanucci 2014). Facing this lack of diagnostic information, morphological studies have struggled to characterise Ischnocolinae relationships. It is hence not surprising that the subfamily as a whole has earned a reputation for being something akin to a “holding place” for theraphosids that cannot be placed into other lineages. Consequently, morphology-based studies of Ischnocolinae recovered the group as paraphyletic. In his broad morphological study, Raven (1985) identified at least four independent clades within Ischnocolinae, with three of them proposed as the earliest branching theraphosids in his phylogeny. Later, Guadanucci (2014) performed phylogenetic analyses on all known genera of Ischnocolinae, plus representatives of the other subfamilies, again based on morphology. He identified the “core group” (Ischnocolinae *sensu stricto*) to include the type genus of the subfamily (*Ischnocolus*) as well as *Acanthopelma, Holothele, Reichlingia* and *Trichopelma* (Guadanucci, 2014). Some other Ischnocolinae (*sensu lato*) were placed in a separate clade, including *Schismatothele* plus others including several former *Holothele* species later transferred to other genera (including *Neoholothele*), as the newly erected subfamily of Schismatothelinae (Guadanucci, 2014; Guadanucci & Weinmann, 2015). Remaining ischnocoline genera, such as *Heterothele, Nesiergus* and *Catumiri*, were dispersed across the phylogeny, and their phylogenetic affinities remained largely unresolved. Our phylogenomic analysis, for the first time, represents a non-morphology based multigene hypothesis for theraphosid subfamily relationships that includes more than one member of Ischnocolinae. We included *Trichopelma laselva* (Ischnocolinae *sensu stricto*) *Neoholothele incei* (a former ischnocoline now in Schismatothelinae), and *Catumiri* sp. (representing Ischnocolinae *sensu lato*). In agreement with Guadanucci (2014), we detected no obvious close phylogenetic affinity between these three “ischnocoline” taxa, and instead found that *Trichopelma laselva* shares a more recent common ancestor with other Neotropical subfamilies, including Aviculariinae and Theraphosinae, than it does with *Catumiri* sp. Interestingly, members of Schismatothelinae were also found inside this “bombardier clade”, nested between Aviculariinae and Psalmopoeinae, in a placement that sufficiently distinguishes them from other sampled “ischnocolines”. In contrast, *Catumiri* sp. was found as one of the earliest branching theraphosids in our tree as a sister group to Eumenophorinae. In the recently published molecular phylogeny of Lüddecke et al. (2018), with Nesiergus insulanus from the Seychelles only a single member of Ischnocolinae was included, which was seemingly most closely related with Asian Selenocosmiinae. Put together, we conclude that, as implied by Guadanucci (2014), the “Ischnocolinae” as currently defined represent multiple independent theraphosid lineages. We emphasize the need for future large scale integrative studies that are inclusive of problematic members from across the tarantula family, with a particular focus including multiple “ischnocoline” from diverse geographical areas, plus several representatives of Schismatothelinae, which has also been shown to likely be paraphyletic (Lüddecke et al., 2018; Hüsser, 2018). Unfortunately, we were unable to include additional Schismatothelinae taxa in this study, and therefore cannot test the monophyly or paraphyly of the subfamily.

Finally, *Ephebopus* is one of the most controversially placed genera of Theraphosidae in recent times worthy of mention. This genus has only recently been recognized as part of the Psalmopoeinae which otherwise consists of the genera *Psalmopoeus, Pseudoclamoris* and *Tapinauchenius* (after Hüsser, 2018). This subfamily is represented in our study by four species: *Ephebopus cyanognathus, Psalmopoeus cambridgei, Psalmopoeus irminia* and *Tapinauchenius violaceus*. Various members of this clade have commonly been considered as members of the Aviculariinae (e.g. Schmidt 2003; West et al., 2008; Fukushima and Bertani, 2017), or Selenocosmiinae (e.g. Guadanucci, 2014), although some were also placed in a separate subfamily, “Psalmopoeinae” by other authors (Samm and Schmidt, 2010). Recently, Lüddecke et al. (2018) and Turner et al. (2018) found molecular evidence for the validity of the subfamily, which shortly thereafter was supported by both morphological and molecular data in Hüsser (2018), Consistent with these studies, our phylogenomic analysis further supports the distinction between Aviculariinae and Psalmopoeinae (including *Ephebopus*), as evidenced by both the high bootstrap support, and by the fact that, once again, Schismatothelinae were found to be closer to Psalmopoeinae than Aviculariinae, which was placed basal to both of these other subfamilies.

### 4.3 Morphology meets Phylogenomics: Insights into the Evolution of Urticating Setae and Defense Strategies in Tarantulas

Based on morphological similarities, and the existence of intermediate forms of urticating setae, Bertani and Guadanucci (2013) proposed that “primitive”, short setae subsequently became modified to urticating setae, and that type III represents an ancestral state that gave rise to types I and IV (see also Pérez-Miles, 2002). This appears to make sense, as types I, III and IV are all found in the subfamily Theraphosinae, and share several morphological similarities — including attachment via a stalk, a distal penetrating tip, and most notably the numerous and sometimes prominent “proximally-facing” barbs along the shaft of the setae. Types II (in Aviculariinae) and V (in *Ephebopus*, Psalmopoeinae) in contrast possess barbs that point distally, but beyond this neither type shows much structural similarity with each other, nor with those in Theraphosinae. This led Bertani and Guadanucci (2013) to conclude that urticating setae evolved at least three times in Theraphosidae, however, their evolutionary history remained speculative. Our data supports the hypothesis that the “proximally-facing” types I, III and IV are homologous (Fig. 3a), and share a common origin in the subfamily Theraphosinae. Further, our ancestral state reconstructions indicate that, as in Bertani and Guadanucci (2013), urticating setae in Theraphosinae, Aviculariinae and *Ephebopus* can each represent an independent origin (multiple-gain scenario, Fig. 3b). However, equally parsimonious alternatives that challenge this hypothesis were recovered. Urticating setae diversity can also be explained by modifications of an ancestral structure that originated once before the divergence of Aviculariinae and Theraphosinae (i.e. earlier than was previously assumed), and then diversified throughout these clades. In this scenario, the trait was subsequently lost twice - once in Schismatothelinae, and again later in some Psalmopoeinae (multiple-loss scenario, Fig. 3c). Another equally parsimonious scenario exists where urticating setae could have again first originated before the divergence of Aviculariinae and Theraphosinae, but were then lost in the ancestor to both Schismatothelinae and Psalmopoeinae when they diverged. Later, urticating setae re-emerged in *Ephebopus* (gain-loss-gain scenario, Fig 3d). Given the structural differences between different types, and the fact that type V are localized to a different area of the body than other types, the first of the three hypotheses (Fig. 3b) seems plausible and is consistent with previous works. However, the other equally parsimonious alternative hypotheses must also be considered

Perafán et al. (2016) described a monotypic genus *Kankuamo* from the Theraphosinae subfamily, which most notably has a novel type of urticating setae (type VII). None of the previously described types displayed much morphological similarity with type II beyond the position criterion (Bertani and Guadanucci, 2013). However, type VII shares several characteristics with type II setae (Perafán et al, 2016). It has even been suggested that urticating setae in *Kankuamo* and Type II share a similar “direct contact” method — i.e. where they are rubbed directly from the opisthosoma onto the target instead of being projected into the air by “bombardment”, though the release mechanism has not been directly observed in *Kankuamo*. Unfortunately, no phylogenetic study has included members of *Kankuamo*, but its placement will likely be crucial in resolving the evolutionary history of urticating hairs. Given their remarkable structural similarity, attachment mechanism and potentially similar method of release, it is plausible that the type II setae in Aviculariinae are derived from type VII in *Kankuamo* or vice versa, providing a link between the type II of Aviculariinae and types I, III & IV in Theraphosinae. This could serve to challenge the prevailing multiple-gain hypothesis presented in Fig. 3b, and potentially lend more support to either of the alternative multiple-loss or gain-loss-gain hypotheses illustrated in Fig. 3c and Fig. 3d respectively.

Type V urticating setae are exclusively present in a single genus, *Ephebopus*, and remarkable in that they are not localised on the opisthosoma, but are instead on the palpal femora (Foelix et al., 2009; Marshall and Uetz 1990b). Both Hüsser (2018) and Bertani and Guadanucci (2013) considered urticating setae to be autapomorphic in *Ephebopus*, inline with our ancestral state reconstruction, although they treated this genus as part of the Aviculariinae. The absence of any urticating setae in Schismatothelinae and the remaining Psalmopoeinae, including our robust placement of *Ephebopus* in that latter family also supports this idea. However, as suggested by Figs. 3b-3d, equally parsimonious alternatives must be considered. Some members of the theraphosine genus *Hemirrhagus* possess type VI urticating setae, which are morphologically very similar to type V (Bertani and Guadanucci, 2013), but are localised on the opisthosoma. *Hemirrhagus* contains many species notable for their unusual troglobitic lifestyles (Mendoza and Francke, 2018). The occupation of subterranean ecological niches, and the switch to a cave-dwelling lifestyle, has probably contributed to the evolution of unique and unusual morphological traits in *Hemirrhagus*. On the other hand, other traits have been lost in some species, including eye pigmentation and urticating setae, which have been interpreted as derived reversals (Mendoza and Francke, 2018). If members of this genus can be included in future studies, it may pave the way to study general patterns of ecological adaptation, adaptive potential and trait evolution in Theraphosidae. Unfortunately, we were unable to obtain members of *Hemirrhagus* for this work, but in our view, the genus represents an interesting taxon that will be vital to consider in further studies on the evolution of tarantulas. We emphasise that future studies specifically addressing the evolution of urticating setae within the bombardier clade should put particular focus on additional insights from both *Kankuamo* and *Hemirrhagus*.

In previous studies, the subfamilies of our newly established “bombardier clade” were scattered across the tarantula phylogeny, or formed unresolved polytomies with others (e.g. Raven 1985; Guadanucci 2014). In our study, for the first time, we recover the majority of New World theraphosids together as a well-supported monophyletic group, which allows us to suggest that the evolution of urticating setae represents a key innovation that could have facilitated their rapid adaptive radiation in the New World. This hypothesis is further supported by the fact that the Theraphosinae subfamily, in which the diversity and abundance of urticating setae is most conspicuous, represents the largest radiation of known genera and species within Theraphosidae. In fact, this subfamily accounts for about 50% of the known theraphosid diversity (Kambas, 2018), with several types of urticating setae (e.g. Schmidt, 2003). In conclusion it appears reasonable to assume that this high success in the New World might be facilitated via the invention of defensive urticating setae. Like all spiders, members of the “bombardier clade” are venomous, and are capable of delivering defensive bites to would-be predators. Generally, venom serves the purposes of subduing prey, deterring competitors or defense against predators (Casewell et al., 2013). However, venom is foremost a costly resource, and any utilization that conserves as much venom as possible should be considered a competitive advantage (Morgenstern and King 2013). Although tarantulas in general are not particularly venomous to vertebrates such as humans, a small fraction of species seems to be capable of delivering medically significant bites. As scientific literature currently only treats members of *Poecilotheria* as potentially dangerous, bites from other species of African and Asian theraphosids have anecdotally been reported to cause relatively severe and painful symptoms (Escoubas and Rash, 2004; Hauke and Herzig, 2018). It is intriguing that all species from which medically significant bites and potent venom are reliably reported lack the defensive mechanism of urticating setae. The study of venom evolution elsewhere has taught us that venom components in ancient lineages, such as spiders, often evolve under strong purifying selection pressures, which predicts that over the course of evolution, only indispensable components remain present in venom cocktails (Sunagar and Moran, 2015). In light of this, it becomes possible to imagine a scenario in which the evolution of inexpensive urticating setae may have led to the loss of costly venom components, possibly as an adaptive response to a new environment.

### 4.4 Contentious Nodes

Despite the overall high support and strong phylogenetic signal in our study, as revealed from maximum bootstrap and high gene jackknife support for most nodes, two nodes were not strongly supported in all analyses and require more detailed discussion.

The first of these corresponds to the *Lasiodora parahybana + Aphonopelma johnnycashi* node, which was not recovered by ASTRAL. In matrices 1-4, *Lasiodora parahybana* was recovered as the sister taxon to *Aphonopelma johnnycashi, Acanthoscurria geniculata* and *Phormictopus atrichomatus*. Matrix 5 (MARE) did not include *Aphonopelma johnnycashi*, but adjacent nodes still recieved 100% bootstrap support. ASTRAL instead placed *Aphonopelma johnnycashi* as sister to *Lasiodora parahybana, Acanthoscurria geniculata* and *Phormictopus atrichomatus*. Although this means the relationship did not receive 100% support across all analyses, the general relationship between the two was recovered across all six analyses, and we are quite confident in the validity of this node based on the strong bootstrap support from matrices 1-5, which each recovered *Lasiodora parahybana* as sister to *Aphonopelma johnnycashi*.

Our second poorly supported node may instead highlight some key limitations of the core-ortholog approach. Our phylogeny shows Selenocosmiinae + Thrigmopoeinae as early branching subfamilies, but they were found to branch alongside Poecilotheriinae / Ornithoctoninae / Harpactirinae / Stromatopelminae in matrices 2 and 3. This topology was not strongly supported in any of our analyses. Prior to the addition of the gene tree pruning step to further minimise paralogy, this node had even weaker support — it was only recovered in 3 of our 6 topologies (as opposed to 4 of 6 post-pruning), with matrix 3 yielding the greatest highest value of 62% for that node despite no longer recovering it post-pruning. We caution future analyses that use a core-ortholog approach that this part of the phylogeny may be particularly susceptible to some influence of paralogous genes, and that careful filtering is warranted. Yet, the poor support of this node remains somewhat of an enigma, especially given that the respective branches are not particularly short, which would otherwise suggest a fast radiation.

Possibly, the support values for this particular node could be improved with a greater breadth of taxon sampling. In particular, we believe that the addition of a member of Selenogyrinae (the only subfamily absent from our phylogeny) could be key to breaking up these long branches and resolving this node. Selenogyrinae are found throughout west / central Africa, as well as India, though the placement of the latter in this subfamily may be questionable. Given the strong biogeographic links we found across the phylogeny, it is reasonable to suppose that Selenogyrinae would emerge alongside other subfamilies found in these regions., but it seems plausible that Selenogyrinae could help resolve this uncertainty based on their geographic spread. Unfortunately, we recognize that these taxa are very difficult to procure, with new field-collection likely to be the only source for future studies.

### 4.5 Future Perspectives

Although our phylogeny provides a robust framework for future evolutionary studies on theraphosids, there are still some potential sources of bias that should be discussed. It has been shown that taxon selection and the density of sampled taxa represent important factors that influence tree topologies, and therefore the inferred hypothesis (e.g. Pollock et al., 2002; Zwickl and Hillis, 2002). Increased taxon coverage from both within Theraphosidae and from other mygalomorph families can benefit our understanding of theraphosid evolution. Relatively little is known about the family Barychelidae, commonly known as brushed-foot trapdoor spiders, who are a sister group to Theraphosidae (Hamilton et al., 2016) and a group for which no transcriptome data is currently available. The relationships between these families are tenuous, and some theraphosids have a long yet recent history of being treated as barychelids — e.g. Guadanucci (2014), a relatively recent publication, determined that many genera, including *Trichopelma* (represented in our study by *T. laselva*), actually belong in Theraphosidae and not Barychelidae as was previously thought. This alone should signal a need to more carefully re-examine the relationships between theraphosids and barychelids using phylogenomic analyses.

Our phylogenomic analysis, despite including representatives of all major theraphosid lineages, is based on data from only 25 of the 146 currently accepted genera (i.e. 18%), and it is apparent that our dataset only captures a subset of the whole radiation of this incredibly species-rich family (World Spider Catalog 2018). It should therefore be a high priority for future studies to expand the inherent taxon coverage within theraphosids to provide a more complete picture of the family’s evolution. We suggest that the expansion of taxon sampling in future research should further be economised and focused on lineages of uncertain taxonomic status, such as Ischnocolinae, Selenogyrinae and Thrigmopoeinae, or taxa from currently neglected geographic areas such as the Indian subcontinent, Europe or Australasia. Moreover, as previously mentioned, the addition of further taxa with unique urticating setae types will be required to capture the full gamut of diversity of this unusual character, which in turn will be invaluable to determine exactly how this trait has evolved. That said, the study of venom evolution, in particular the relationship between toxins and urticating setae possession, might be possible to evaluate in the future in the context of our robust phylogenomic reconstruction, especially should more taxa from the above mentioned groups be included.

## 5. Conclusion

Our research provides the first phylogenomic hypothesis on the evolution of tarantula spiders, and helps clarify possible evolutionary scenarios pertaining to their urticating setae. Our phylogeny differs from a wide variety of previously published morphological and molecular studies. Deeper nodes illustrating the relationships between subfamilial lineages have been recovered with strong support for the first time on a robust phylogeny, which can serve as a sturdy foundation for a diverse range of subsequent studies on these enigmatic creatures.

## 6. Acknowledgements

We are grateful to the German Arachnological Society (Deutsche Arachnologische Gesellschaft, DeArGe e.V.) for providing funds to this study, and to the National Parks Board of Singapore for permission to collect *Phlogiellus inermis* (permit number NP/RP18-046). We thank Iker Irrisari for generating the initial jackknife replicates, and David Court and Björn Marcus von Reumont for valuable discussions on the manuscript. Bastian Rast kindly provided picture material that has been used to illustrate the phylogenetic tree. We are further thankful to Martin Hüsser and Rainer Foelix for providing illustrations that were used to assemble our figure on urticating setae.

## 7. Author Contributions

SF and TL designed the study and acquired sample material. SF, TL and SK performed laboratory work. TL, MV and WHP attracted funding for the study. Bioinformatic work was conducted by SF, WHP, MV and DQC. HK, SJL, IW, VvW and RT contributed to experimental design and assured valid taxonomical as well as morphological assignments. SF and TL wrote the manuscript with substantial input from MV, HK, SJL, IW, VvW and RT. All authors reviewed and agreed on the manuscript.

